# The central nervous system is a potential reservoir and possible origin of drug resistance in HBV

**DOI:** 10.1101/2023.04.05.535804

**Authors:** Lijun Xu, Minghan Zhou, Xiuming Peng, Yufan Xu, Fan Huang, Linyun Wang, Xiaorong Peng, Zongxing Yang, Ran Tao, Guanjing Lang, Qing Cao, Minwei Li, Ying Huang, Biao Zhu, Yan Xu

**Affiliations:** Department of Infectious Diseases, the First Affiliated Hospital, College of Medicine, Zhejiang University, Hangzhou, China; National Clinical Research Center for Infectious Diseases, the First Affiliated Hospital, College of Medicine, Zhejiang University, Hangzhou, China; The State Key Laboratory for Diagnosis and Treatment of Infectious Diseases; the First Affiliated Hospital, College of Medicine, Zhejiang University, Hangzhou, China; Department of respiratory medicine, Sir Run Run Shaw Hospital, College of Medicine, Zhejiang University, Hangzhou, China; Department of Pathology, Sir Run Run Shaw Hospital, College of Medicine, Zhejiang University, Hangzhou, China; Department of Infectious disease, Jianxin Hospital, Fuzhou, China; College of Medicine, Zhejiang University, Hangzhou, China; Department II of Infectious diseases, Xixi hospital, College of Medicine, Zhejiang University, Hangzhou, China

**Keywords:** HBV, cerebrospinal fluid, sequence, mutation

## Abstract

The significance of HBV in cerebrospinal fluid (CSF) is unclear. In the present study, **s**ynchronous serum and CSF samples were collected from 13 patients. HBV-DNA, full-length genome, quasispecies, phylogenetic tree, compartmentalization and mutation of reverse transcriptase (RT) region analyses were performed based on PCR and sequencing methods. We found HBV-DNA was detected in the CSF of 3 antiviral-naïve patients and one patient after successful antiviral therapy. Complete full-length HBV genomes were isolated from the CSF of 5 patients, including 2 patients with undetectable serum HBV-DNA. Ten patients exhibited distinct CSF-serum quasispecies, 8 harbored independent CSF-serum genetic compartmentalization and phylogenetic trees, and 5 patients presented lamivudine/entecavir-associated resistance mutations in only the CSF. The frequencies of rtL180M and rtM204I/V mutations in both serum and CSF were higher in HIV/HBV-coinfected patients than in HBV-monoinfected patients (serum: rtL180M: 3.9% vs. 0, P = 0.004; rtM204I/V: 21.3% vs. 0, P < 0.001; CSF: rtL180M: 7.6% vs. 0, P = 0.026; rtM204I/V 7.6% vs. 1.6%, P = 0.097). Our data suggested CSF is a potential HBV reservoir, and HBV in CSF harbors distinct evolution and mutation models from that in serum. HIV infection increases the possibility of HBV rtL180M and rtM204I/V mutations in both serum and CSF.

## INTRODUCTION

Whether the central nervous system (CNS) is an HBV reservoir is currently controversial. Some studies have demonstrated that hepatitis B surface antigen (HBsAg) and hepatitis B core antigen (HBcAg) are detectable in brain tissues upon autopsy^[1]^, which raises an interesting question: is HBV-DNA present in cerebrospinal fluid (CSF)? It is commonly believed that HBV cannot cross the blood‒brain barrier (BBB)^[2]^. However, some studies have found faint traces of HBV sequences and markers in patients suffering from neurological disorders, such as fatal Creutzfeldt‒ Jakob disease, cryptococcal meningitis and other neurological diseases^[3–5]^. In addition, next-generation sequencing (NGS) analysis has revealed HBV-DNA sequences in the CSF of patients with viral encephalitis^[6]^. These data indicate that HBV-DNA sequences could be present in the CSF of patients with some CNS disorders. However, whether the full-length HBV genome exists in the CNS remains unknown.

Lymphocytes and neutrophils are usually poorly recruited in normal CSF. Therefore, the CNS might be an ideal refuge for HBV because of its low immune response. If so, are HBV quasispecies in CSF different from quasispecies in serum? Complex quasispecies of HBV in extrahepatic reservoirs might be related to drug resistance and recurrence of HBV^[7–9]^. Of note, the currently used anti-HBV drugs, such as lamivudine (LAM or 3TC) and tenofovir disoproxil fumarate (TDF), have poor BBB penetration activity^[10, 11]^. Presumably, low immune pressure and drug pressure in CSF probably have potential effects on the replication and evolution of HBV in the CNS and even on drug resistance.

In the present study, 13 HBV-infected patients were recruited to compare the structure of the full-length genome, compartmentalization, quasispecies and reverse transcriptase (RT) region variants of HBV between serum and CSF samples with the aim of elucidating the implication of CSF HBV in clinical practice.

## MATERIALS AND METHODS

### Patients

Thirteen participants (10 HIV/HBV-coinfected and 3 HBV-monoinfected) were recruited for this study: 3 patients with cryptococcal meningitis, 2 patients with neurosyphilis, one patient with *Toxoplasma gondii* infection, and 7 patients with a suspected CNS disorder who underwent lumbar puncture but were not confirmed to have CNS disease. The clinical indication for lumbar puncture was determined by physicians according to the following criteria: (1) routine lumbar puncture for patients with confirmed CNS diseases (e.g., cryptococcal meningitis and syphilis) and (2) lumbar puncture for patients with presumable illnesses in the CNS (e.g., cerebrovascular disease or nonspecific headaches). Synchronous CSF and serum samples were obtained after participant authorization. The samples were collected between Jan 2014 and Dec 2018 and frozen at −80°C for subsequent analysis.

### Extraction of HBV-DNA

Serum and CSF samples were collected on the same day as the lumbar puncture. HBV-DNA was extracted from 200 μL of serum/CSF samples using a QIAamp Blood Mini Kit (Qiagen, Hilden, Germany) according to the manufacturer’s instructions, and the purified DNA was dissolved in 40 μL of distilled water for further use.

### Quantitation of HBV-DNA in serum and CSF

Quantitative PCR (qPCR) was used to quantify the HBV-DNA levels. A total of 600 µL of serum or CSF was used for qPCR according to the standard protocol of the COMBAS TaqMan 96 HBV Test V2.0 Kit (Roche, IN, USA), and qPCR was performed with the Roche COBAS Ampliprep PCR system (Roche, IN, USA). The detection range was from 20 to 1.7◊10^8^ IU/mL.

### Isolation of full-length HBV genomes in blood and CSF

Nested PCR was used to amplify the full-length HBV genome. The PCR primers and conditions were set according to Gunther’s method^[12]^. After two rounds of PCR amplification, a full-length 3.2-kb or another approximately 2.0-kb PCR product of the HBV genome was obtained.

The PCR products (3.2 kb or 2.0 kb of HBV-DNA if the 3.2-kb DNA was unavailable) were cloned into the vector PMD-19T with T4 DNA Ligase (Takara, Japan) and transformed into TOP10 Escherichia coli competent cells (Invitrogen, CA, USA). Transformants were grown on ampicillin-supplemented plates. Positive clones were identified by PCR. Approximately 1-2 randomly chosen clones per sample were used for the identification of HBV genomes.

### Cloning and sequencing of the HBV RT region

The HBV RT region was amplified by nested PCR with two-round PCR primers according to the method described by Chen et al^[13]^. After two rounds of amplification, an approximately 1100-bp fragment encompassing the entire RT region was obtained, cloned into the PMD-19T vector (Takara, Japan) and then transformed into TOP10 *Escherichia coli* competent cells (Invitrogen, USA). The transformants were grown on ampicillin-supplemented plates. Approximately 20 randomly chosen clones per sample (one patient had only 8 clones in the CSF sample) were sequenced and analyzed.

### Analysis of HBV quasispecies based on the RT region

Multiple sequence alignment was performed using CLUSTAL X (version 2.0). The HBV quasispecies were evaluated for complexity and diversity according to previously described methods ^[13]^. The quasispecies complexity, known as Shannon entropy (Sn), was calculated using the following formula: Sn = −∑_i_ (pi lnpi)/lnN (N is the total number of clones, and pi is the frequency of each clone in the viral quasispecies population). The quasispecies complexity was calculated at the nucleotide and amino acid levels. Sn ranges from 0 to 1 (0 means that all virus sequences are identical, and 1 means that each sequence is different and unique^[14]^). The quasispecies diversity was calculated for each patient by the mean genetic distance (d), including the number of synonymous substitutions per synonymous site (dS) and the number of nonsynonymous substitutions per nonsynonymous site (dN). The genetic distances at the nucleotide level were calculated under the Tamura three-parameter model that incorporates transitional and transversional rates and G + C content bias, whereas the genetic distances at the amino acid level were calculated under the JTT model. The dS and dN were calculated under the modified Nei– Gojobori model with Jukes–Cantor correction. Phylogenetic trees were constructed with the maximum likelihood method under the Tamura-Nei model. Mega11 software and the online tool https://itol.embl.de/ were used to draw the phylogenetic trees.

### HBV genetic compartmentalization tests

Compartmentalization analysis of serum and CSF samples from each patient was performed using HyPhy software (V2.5)^[15]^. As different methods to define genetic compartmentalization (such as genetic heterogeneity and distinct lineages between two viral subpopulations) might occasionally yield discordant results on compartmentalization tests^[16]^, we applied genetic distance-based tests and tree-based tests for the genetic compartmentalization analysis. The distance-based tests included Fst and Snn (nearest-neighbor statistic), and Fst has several different types: 1) Hudson, Slatkin and Madison (HSM), 2) Slatkin (S), and 3) Hudson, Boos and Kaplan (HBK). We employed the HBK nonparametric test for the distance-based test and computed the *K_ST_* statistic, whose values range from 0 (denoting no compartmentalization) to 1 (denoting complete compartmentalization). In addition, the Slatkin-Maddison (SM) method was used for the tree-based test. A p lower than 0.05 indicates that the elements can be judged to be compartmentalized. Compartmentalization was finally confirmed by two algorithms (*K_ST_* and SM tests) when both p values were less than 0.05.

### Analysis of drug resistance in the HBV RT region

Drug resistance to lamivudine (LAM, 3TC), entecavir (ETV) and tenofovir disoproxil fumarate (TDF) was defined as in previous reports^[17–19]^:

LAM (3TC) resistance was defined by detection of any mutation within the RT region as follows: rtL80I, rtL173M, rtL180M, and rtM204I/V.

ETV resistance was defined by detection of the combination of rtL180M and rtM204I/V mutations (rtL180M + rtM204I/V) plus any of the following mutations: rtL80I, rtI169V/M, rtA181G/C, rtT184A/F/I/L/M/S, rtS202G and rtM250V.

TDF resistance was defined by detection of four mutations, namely, rtS106, rtH126, rtD134 and rtL269.

An online tool (https://hivdb.stanford.edu/HBV/HBVseq/development/HBVseq.html) <u>was us</u>ed to analyze the mutations in the RT region and genotypes of HBV. A graphical representation of the amino acid sequence alignment of the HBV RT region was drawn using an online tool (https://weblogo.threeplusone.com). Each logo consists of stacks of symbols, with one stack for each position in the sequence. The overall height of the stack indicates the sequence conservation at that position, whereas the height of symbols within the stack indicates the relative frequency of each amino at that position.

### Statistical analysis

Continuous normally distributed variables are presented as the means ± standard deviations, and continuous nonnormally distributed variables are presented as medians (interquartile ranges, IQRs). Categorical variables are presented as the numbers of cases (percentages). Continuous variables were compared by Student’s t test or the Mann‒Whitney U test, whereas categorical variables were compared using the χ2 test or Fisher’s exact test. A P value <0.05 (two-tailed) was considered to indicate significance. The data analyses were performed using SPSS version 24.0 (IBM, NY, USA).

### Ethics

Written consent was obtained from all the patients. The study was in accordance with the 1975 Declaration of Helsinki and approved by the Ethics Committee of the First Affiliated Hospital, School of Medicine, Zhejiang University (No. 2014-335). All patient data were anonymously analyzed.

## RESULTS

### Basic demographic characteristics of patients

The demographic details, morbidities and initial CSF profiles of the patients are listed in **Table 1**. The average age was 39.7± 10.7 years. Three patients accepted anti-HBV drugs (HBV-HIV-coinfected patient 4 and 8 accepted ≥2-month anti-HIV regimen of TDF+3TC+efavirenz, and HBV-monoinfected patient 3 accepted the 1-year ETV treatment) at enrollment. Among 11 patients with sufficient serum samples and 8 patients with sufficient CSF samples, serum HBV-DNA was detectable in 8 patents with a mean level of 6.5±1.9 (lg) IU/mL, and CSF HBV-DNA was detectable in 4 patients with a level of 2.4±0.7 (lg) IU/mL. There were 7 patients with genotype B HBV and 4 patients with genotype C HBV in serum, whereas there were 6 patients with genotype B HBV and 5 patients with genotype C HBV in CSF. Notably, 2 patients (patients 1 and 2) exhibited discordant genotypes between serum and CSF.

**Table 1.**
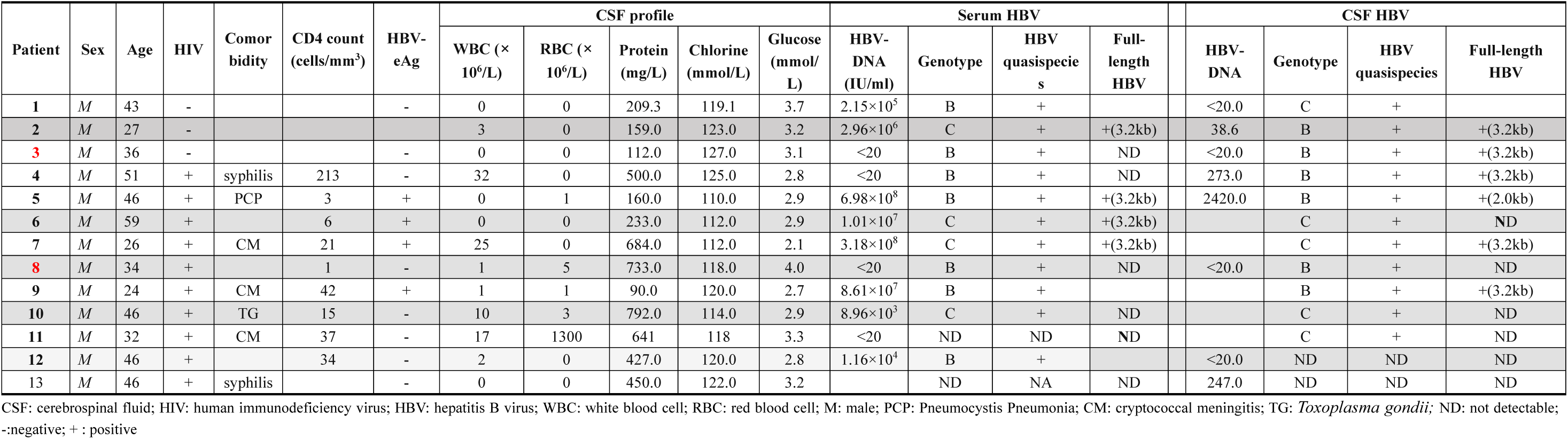
Basic information of HBV-infected patients.

### Full-length HBV genomes were detectable in CSF samples

The full-length HBV genome was successfully isolated from serum samples of 4 patients and from CSF samples of 5 patients. In addition, a 2.0-kb HBV genome fragment was isolated from the CSF of patient 5. Of those, HBV genomes were isolated from both the serum and CSF samples of 3 patients (2, 5 and 7) (**Table 1**). Notably, the full-length HBV genome was obtained in the CSF of patient 3 and 4, although their serum HBV-DNA level were < 20 IU/mL.

A map of the HBV genome structure was drawn according to the full-length sequence of HBV. PreS1 and S2 region deletion mutations in the CSF HBV genome were found in 3 patients (3, 4 and 9). Patient 5 had PreS1 and S2 region deletion mutations in serum but only PreS2 deletions in CSF, whereas patient 6 had a partial gene deletion in Pre S2 in serum (**Figure 1**).

**Figure 1.**
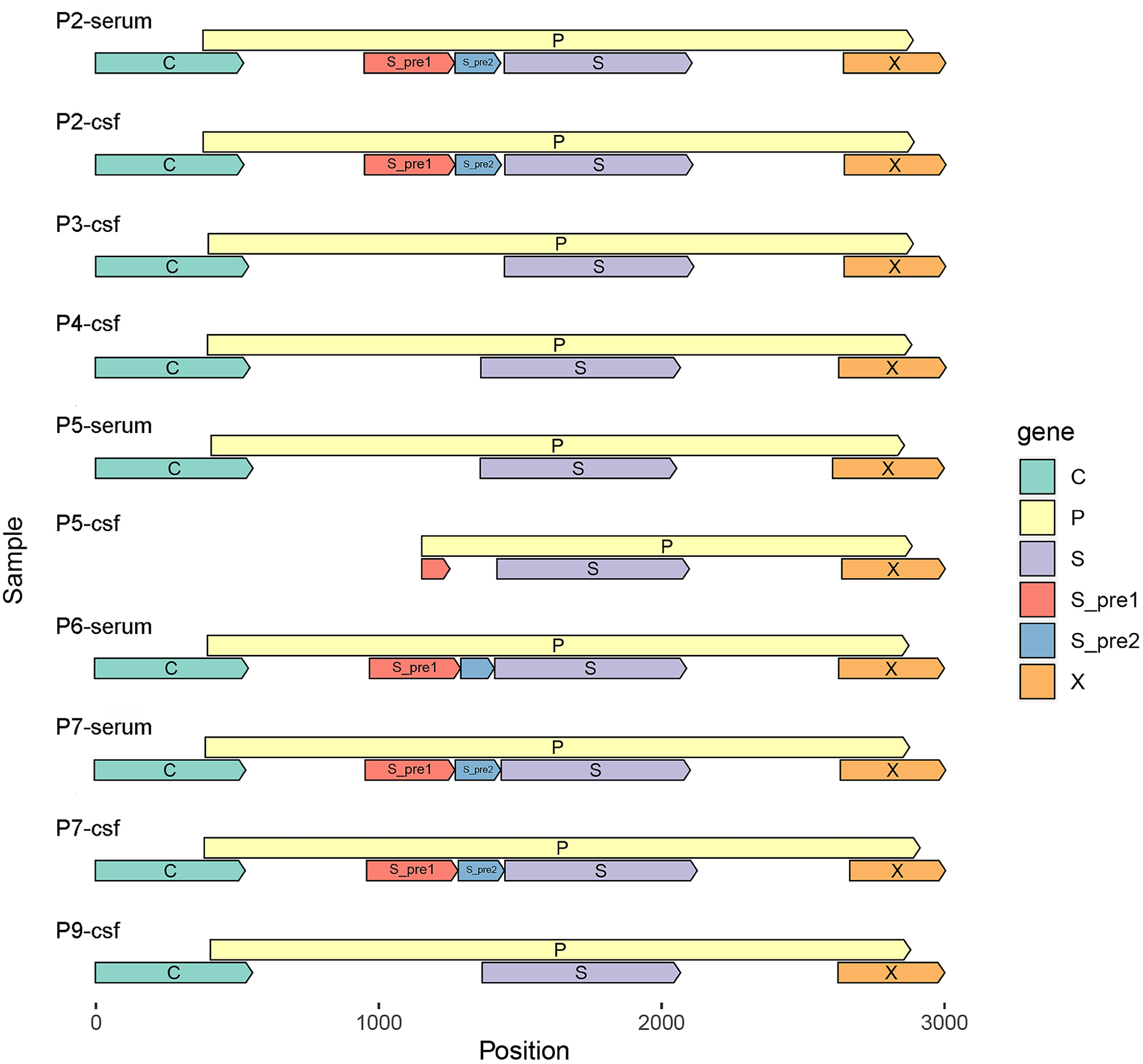
Map of the HBV genome structure. Pre-S1 and S2 region deletion mutations in the CSF HBV genome were found in patients 3, 4 and 9. Pre S1 and S2 region deletion mutations were detected in the serum of patient 5, but only Pre S2 deletion mutations were found in the CSF of this patient 5. P1-P10: patient 1-patient 10.

### CSF HBV quasispecies were divergently distinct from serum HBV quasispecies

Serum HBV sequences of the RT region from 11 of 13 patients andCSF HBV sequences of the RT region from 11 of 13 patients were obtained, respectively. Of those, simultaneous CSF and serum HBV sequences of the RT region were isolated from 10 patients. Notably, HBV RT region sequences were also isolated from CSF samples of patients 3 and 11, although their serum HBV-DNA levels were less than 20 IU/ml (Table 1).

The HBV quasispecies complexity ranged from 0.944 to 1.000 in serum and 0.943 to 1.000 in CSF at the nucleotide level and from 0.913 to 1.000 in serum and 0.865 to 1.000 in CSF at the amino acid level, indicating that all HBV sequences in CSF were highly complex and unique compared with the sequences in serum. Furthermore, the composition and distribution of HBV quasispecies in CSF were also obviously divergent from the quasispecies in serum (**Table 2** and **Figure 2**).

**Figure 2.**
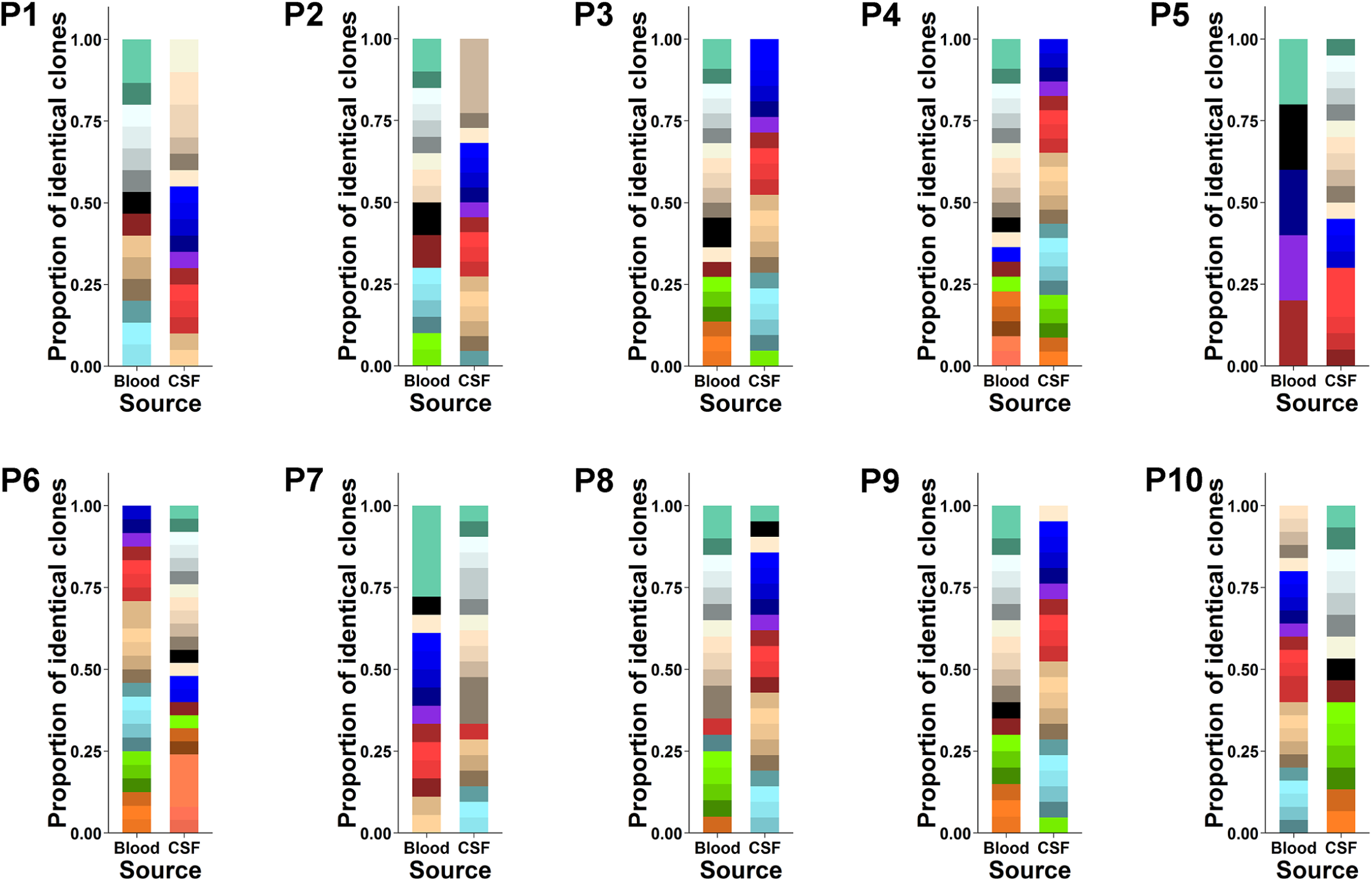
Composition and distribution of quasispecies in serum and CSF samples. Each color represents one kind of quasispecies. The vertical bars indicate the number and proportion of viral quasispecies within each sample. Complex and unique HBV quasispecies were found independently in serum and CSF, respectively. P1-P10: patient 1-patient 10.

**Table 2.** Complexity of HBV quasispecies at nucleotide level and amino acid level.

| patient | Nucleotide level |  | Amino acid level |  |
| --- | --- | --- | --- | --- |
|  | serum | CSF | serum | CSF |
| 1 | 0.991 | 0.984 | 0.991 | 0.865 |
| 2 | 0.984 | 0.943 | 0.984 | 0.928 |
| 3 | 0.990 | 0.994 | 0.913 | 0.994 |
| 4 | 0.995 | 1.000 | 0.978 | 0.979 |
| 5 | 1.000 | 0.979 | 1.000 | 0.975 |
| 6 | 0.994 | 0.984 | 0.966 | 0.971 |
| 7 | 0.944 | 0.982 | 0.922 | 0.962 |
| 8 | 0.989 | 1.000 | 0.971 | 0.994 |
| 9 | 0.994 | 1.000 | 0.994 | 1.000 |
| 10 | 0.998 | 0.997 | 0.996 | 0.974 |

The quasispecies diversity, including d, dS and dN, was compared between serum and CSF samples of host patients. We found that patients 1, 2, 4 and 5 had higher d, dS and dN values for the HBV RT region in serum than in CSF (P < 0.005). Moreover, patient 3 had lower d, dS and dN values in serum than in CSF (P < 0.005), patients 6 and 9 had lower d and dS values (P < 0.05) in serum but a similar dN value in CSF (P = 0.150 and P = 0.801, respectively), and patient 7 had lower d and dN values in serum (P < 0.005) but a similar dS value in CSF (P = 1.000) (**Figure 3**).

**Figure 3.**
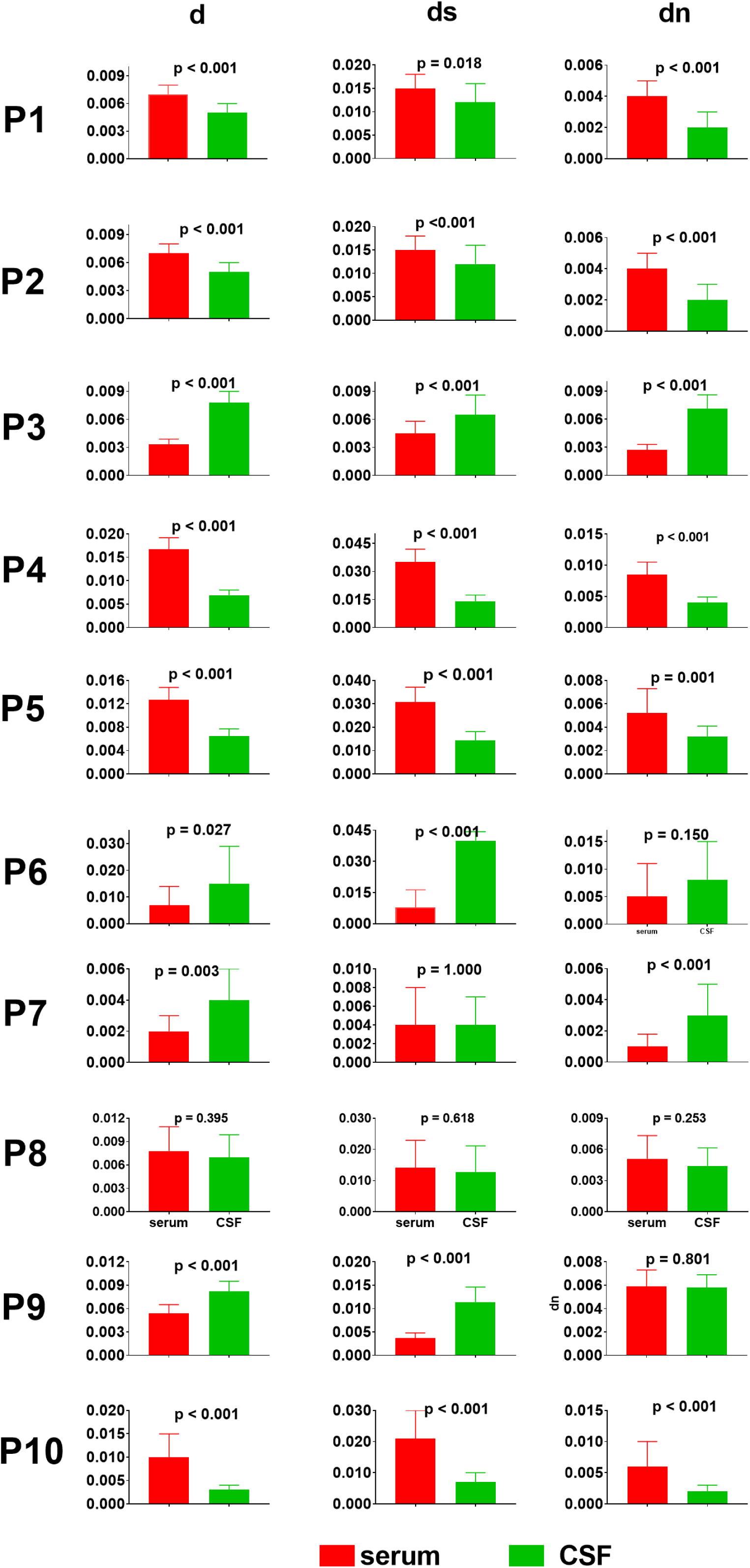
HBV quasispecies diversity in serum and CSF samples. The viral genetic distance (d), synonymous substitutions per synonymous site (dS) and nonsynonymous substitutions per nonsynonymous site (dN) were compared between serum and CSF. P1-P10: patient 1-patient 10.

### The HBV sequences in CSF are highly compartmentalized

The genetic compartmentalization of CSF and serum HBV sequences in 10 patients was analyzed by distance-based tests and tree-based tests. We found that both distance and tree tests agreed for 8/10 patients (including 3/3 HBV-monoinfected patients and 5/7 HIV-HBV-coinfected patients) but not 2/10 patients (patient 5 and patient 8). For patient 5, the distance-based test demonstrated compartmentalization, but the tree-based test did not. Neither tree-based nor distance-based tests revealed blood-CSF HBV genetic compartmentalization in patient 8 (**Table 3**).

**Table 3.**
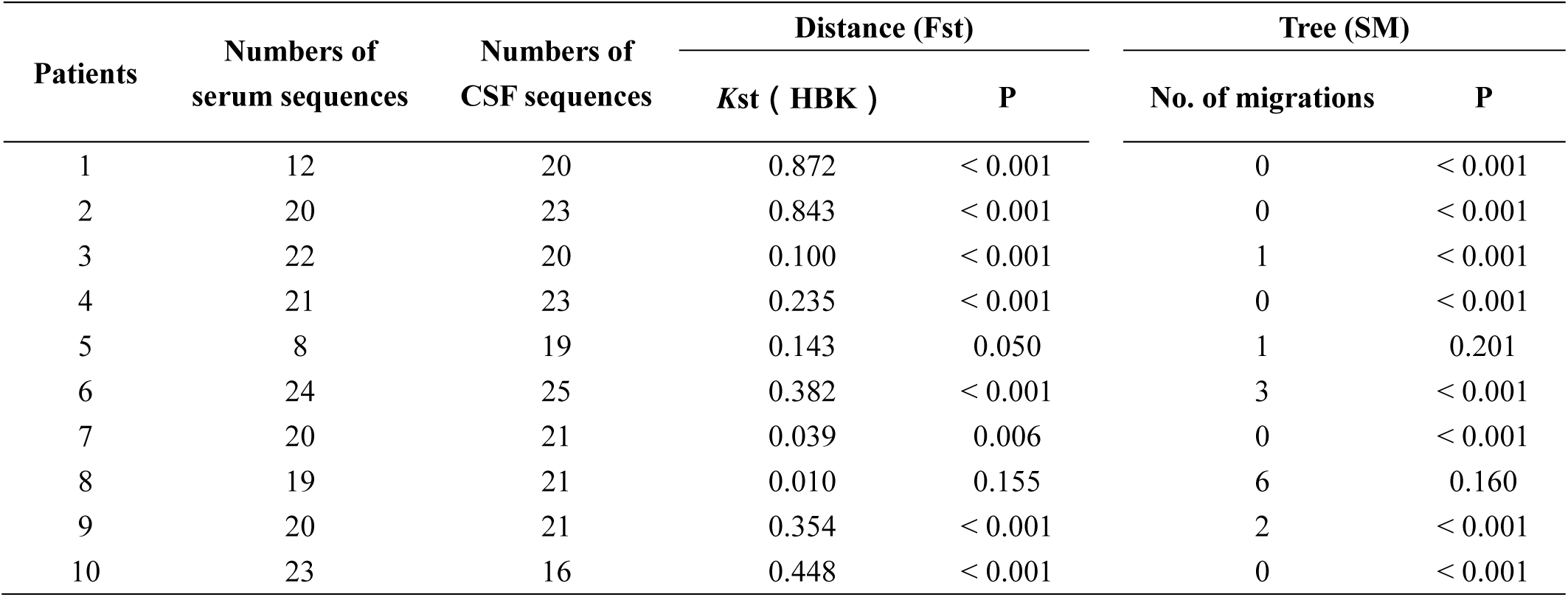
In-host serum-CSF HBV genetic compartmentalization results.

The source of CSF HBV sequences was further analyzed based on RT region sequences. We found that 5 of 23 CSF sequences were amplified from the same clone in patient 2, and one CSF sequence and 5 serum sequences were from one clone in patient 7. More importantly, no clonally amplified sequence was found in the serum or CSF compartmentalized viral population among other patients.

### Phylogenetic analysis of HBV-DNA sequences in paired serum and CSF samples

The phylogenetic trees were constructed under the maximum likelihood algorithm for paired serum and CSF samples in 10 patients. The representative sequences for each patient were used to draw a phylogenetic tree after sequences identified as the same were discarded. We found that 3 patients (1, 2 and 10) harbored highly clustered clades of CSF HBV sequences that were completely separated from clades of serum HBV sequences, and 5 patients (3, 4, 6, 7 and 9) harbored relatively independent clustered clades of CSF HBV sequences separated from serum HBV sequences. Clades of CSF sequences were mixed with clades of serum sequences in 2 patients (5 and 8) (**Figure 4**).

**Figure 4.**
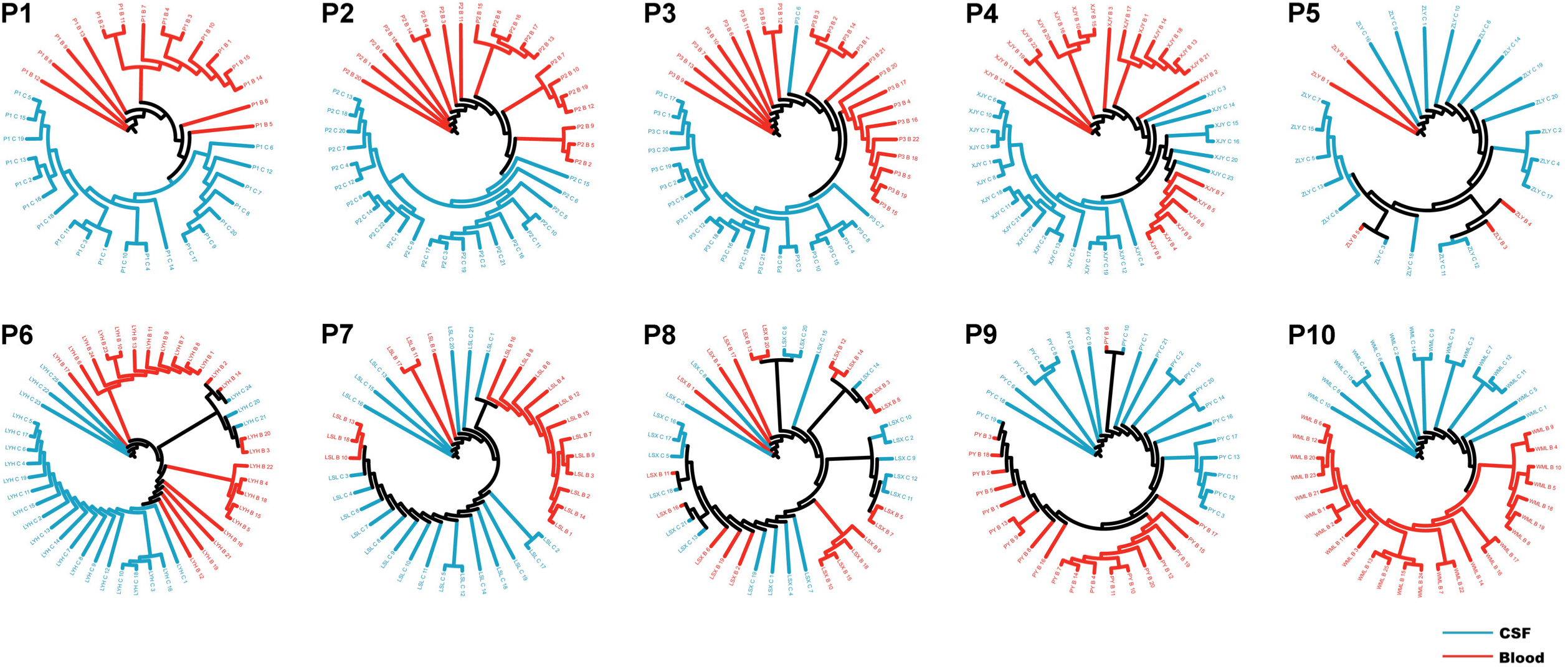
Phylogenetic trees of HBV-DNA sequences in paired serum and CSF samples. The completely or almost clustered clades obtained for the CSF were separated from the clades obtained for the serum samples of patients 1-4, 6, 7, 9 and 10. P1-P10: patient 1-patient 10.

### Drug resistance mutations within the RT region in serum and CSF samples

Drug resistance mutations of LAM (3TC), ETV and TDF within the RT region were analyzed in paired serum-CSF samples.

There were 8.3% (1/12) clones from serum with the rtS202R mutation and 5.0% (1/20) clones from CSF with the rtM204V mutation in patient 1. Sequencing showed 35.0% (7/20) serum clones with rtV84I, 10.0% (2/20) with rtH126D and 75.0% (15/20) with rtD134G/E/S mutations in patient 2. There were 4.5% (1/22) serum clones bearing the rtS202G mutation in patient 3. For patient 4, 4.8% (1/21) of serum clones bore rtV84A, 8.7% (2/23) of CSF clones harbored N236S/D, and 4.3% (1/23) of CSF clones bore rtD134E. The rtD134G mutation [10.5%, 2/19] and rtI169T/M [10.5%, 2/19] were observed in 19 CSF clones from patient 5. For patient 6, triple mutations of ETV resistance in serum clones [95.8% (23/24) rtM204I, 62.5% (15/24) rtM180L and 100% (24/24) rtL80I] and quadruple mutations of ETV resistance in CSF clones [28.0% (7/25) rtM204I, 32.0% (8/25) rtM108L, 28.0% (7/25) rtL80I, 4.0% (1/25) rtM250T] were detected. In addition, patient 6 displayed 4% rtS106F and 4% rtH126R mutations in CSF clones. The rtQ215R mutation (28.0%, 7/25) in serum clones and the rtI169V mutation (4.8%, 1/21) in CSF clones were found in patient 7. Regarding patient 8, rtS106C (47.4%, 9/19), rtD134N6 (8.4%, 13/19) and L269I (100%, 19/19) mutations in serum clones were associated with TDF resistance. Strikingly, quadruple mutations of TDF resistance [rtS106C; 33.3% (7/21), rtH126R: 4.8% (1/21), D134N: 57.1% (12/21), 100% L269I] were found in CSF sequences. Furthermore, the rtT184A and rtS202G mutations were present only in 4.8% (1/21) and 4.8% (1/21) of the CSF clones, respectively. Patient 9 had three TDF resistance mutations (5.0% rtS106P, 5.0% rtD134N and 100% rtL269I) and one ETV-related mutation (rtI169M, 5%) in 20 serum clones. In contrast, 4.8% rtD134N, 100% rtL269I, 4.8% rtV173M and 4.8% rtS202G were found in 21 CSF clones. The rtS202G mutation was found in 4.3% (1/23) of serum clones from patient 10 (**Figure 5**). In addition, the rtL269I mutation was found in all serum sequences and all CSF sequences of patients 3, 4 and 5. Overall, LAM/ETV resistance-associated mutations in only CSF clones were found in 5 patients (patient 1 with rtM204I, patient 6 with rtM250T, patient 7 with rt169V, patient 8 with rtT184A and rtS202G, and patient 9 with rtV173M and rtS202G).

**Figure 5.**
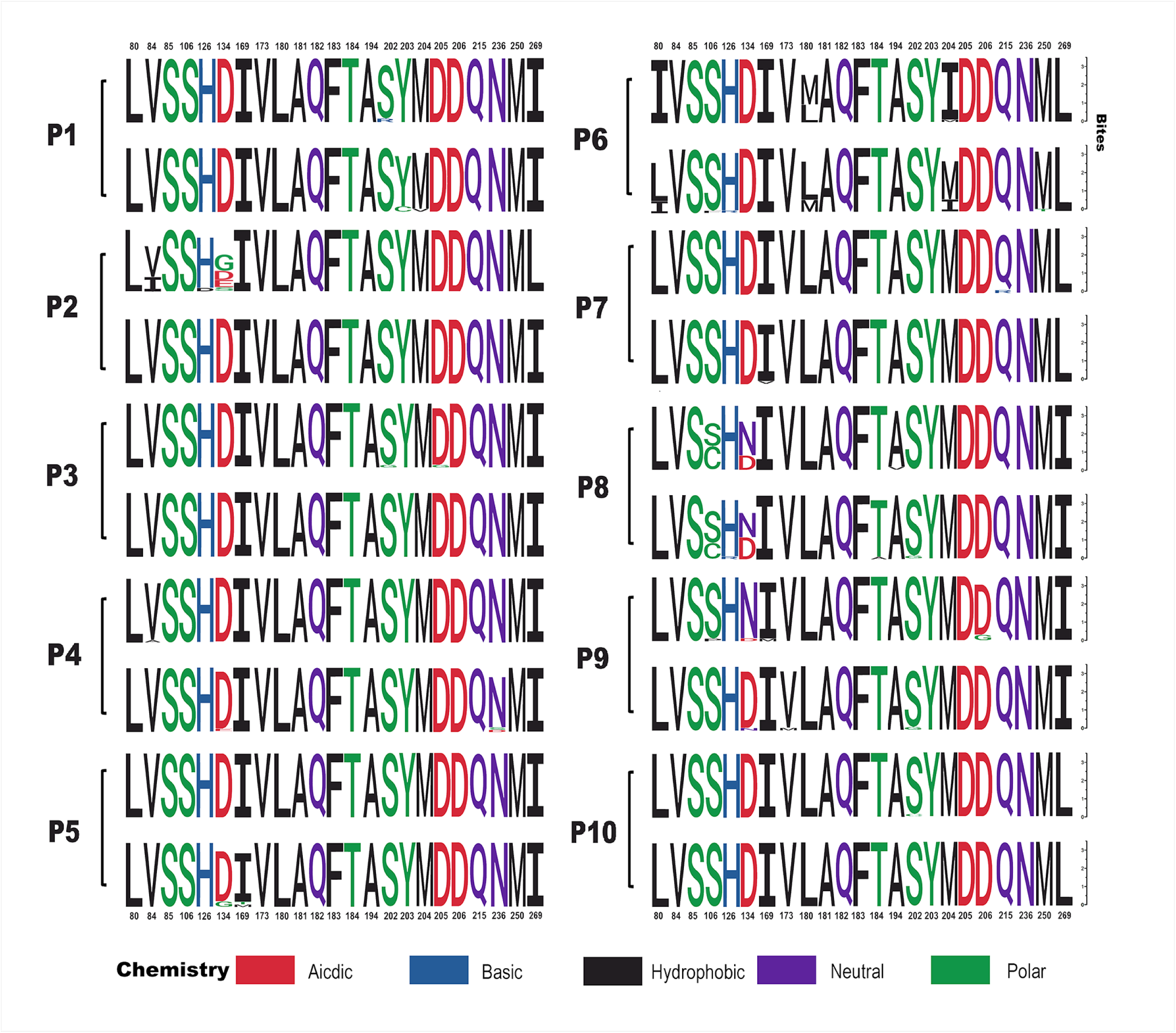
Comparison of major mutations in the HBV RT region between serum and CSF. Serum amino acids are in the upper row, and CSF amino acids are in the lower row. Each logo represents an amino acid sequence. The height of symbols within the stack indicates the relative frequency of each amino at that position.

### HIV infection increased the likelihood of rtL180M and rtM204V/I mutations in both serum and CSF clones

High frequency mutation sites of rtL180M and rtM204V/I were compared between HIV/HBV-coinfected and HBV-monoinfected patients. After patients 5 and 8 were excluded for possible blood contamination of CSF, the rtL180M mutation was found in 13.9% (15/108) of serum clones from HIV/HBV-coinfected patients, but the serum clones from HBV-monoinfected patients did not harbor this mutation (P = 0.004); in addition, the rtL180M mutation was found in 7.6% (8/105) of the CSF clones from HIV/HBV-coinfected patients, and the CSF clones from HBV-monoinfected patients did not have this mutation (P = 0.026). Similarly, the rtM204I/V mutation was found in 21.3% (23/108) of serum clones from HIV-HBV coinfected patients and none of those from HBV-monoinfected patients (P < 0.001), and the rtM204I/V mutation was detected in 7.6% (8/105) of the CSF clones from HIV/HBV-coinfected patients and 1.6% (1/62) of those from the HBV-monoinfected patients (P = 0.097).

## DISCUSSION

Currently, few studies have investigated the evolution and mutations of HBV in the CNS. Our research suggested that 1) full-length HBV genomes could be detectable in the CSF of patients even in those with undetectable serum HBV-DNA; 2) HBV quasispecies in CSF were completely different from quasispecies in serum in terms of both complexity and diversity; 3) the milieu of the CNS fosters unique HBV compartmentalization and phylogenetic trees, indicating an independent evolution model of HBV; and 4) the development of nucleos(t)ide analog resistance mutations in the CNS is probably associated with the presentation of drug resistance in serum.

A previous study indicated that 69.2% (18/26) of HIV/HBV-coinfected adolescents with positive plasma HBV-DNA also have positive CSF HBV-DNA ^[20]^. Another study indicated that the CSF HBV-DNA sequence was detected in two HBV mono-infected patients with CNS disease^[5]^. The full-length HBV genome was found in the CSF of a patient with acute hepatitis B and transverse myelitis^[21]^. In contrast to those studies, our data revealed that HBV-DNA and full-length HBV genomes could be detected in the CSF of adult patients, even those with successful antiviral therapy. These data suggested that the CNS is a reservoir of HBV. Nucleos(t)ide analogs are widely used for HIV and HBV treatment. However, most nucleos(t)ide analogs have poor penetration activity through the BBB^[10, 22]^, and this feature leads to insufficient drug concentrations for HBV clearance in the CNS, which might be related to the persistence of HBV in the CNS. Thus, drugs with higher penetration through the BBB should be considered a new strategy for HBV clearance in the CNS.

Moreover, PreS1 and PreS2 deletions in the CSF HBV genome were detected in our study. It has been reported that the prevalence of PreS deletions in HBV/HIV-coinfected patients is higher than that in HBV-monoinfected patients^[23]^, and a high prevalence of PreS deletion/mutation is associated with immune evasion, advanced liver disease and hepatocellular carcinoma^[24, 25]^. The incidence of PreS1/S2 deletion in CSF samples might reflect the low immune pressure in CSF.

Our data support the notion that the HBV evolutionary profile in CSF is completely different from that in serum based on the following results: 1) the HBV quasispecies were completely different and unique in terms of complexity and diversity between the serum and CSF of most patients; 2) the composition and distribution of HBV quasispecies in CSF were also divergent from those in serum; and 3) the clades of the phylogenetic trees obtained for the CSF were clearly separated from the clades of those found for serum, and this finding was observed for most patients. The distinct HBV population between CSF and serum raises an interesting question: where does the HBV population in CSF originate from? Studies on HIV-1 in the CNS have described three states of the HIV-1 population in CSF compared to that in the blood: 1) equilibrated (when viruses in the blood and CSF are very similar), 2) compartmentalized (when blood and CSF viral populations are distinct, indicating separately evolving populations in these compartments) and 3) clonal amplification (when a single variant is greatly expanded within a compartment)^[26, 27]^. Although the presence of red blood cells in the CSF samples obtained from patients 5, 8, 9 and 10 indicated the possibility of blood contamination of CSF during lumbar puncture, our study illustrated that CSF HBV populations were highly compartmentalized variants from the serum HBV population in most patients except patients 5 and 8. Indeed, the phylogenetic analysis also showed that the CSF clades were independent from the serum clades of most patients. Notably, we also found that one HBV sequence in CSF and five HBV sequences in serum from patient 7 were identical clones, suggesting possible exchange of the HBV between the CSF and serum compartments.

Our data suggested that HBV RT region mutations in the CNS might be a source of HBV drug resistance in peripheral blood. We found that some mutations were only detected in CSF (such as rtM204V in patient 1, rtN236S/D and rtD134E in patient 4, D134G and rtI169T/M in patient 5, and rtS106F, rtH126R and rtM250T in patient 6). Combined mutations conferring ETV resistance in the CSF sequences of patient 6 and quadruple mutations conferring TDF resistance in the CSF sequences of patient 8 were also found in our study (although the HBV sequences in CSF were not compartmentalized). Furthermore, we found that HIV-HBV-coinfected patients had higher frequencies of rtL180M and rtM204I/V mutations in both serum and CSF than HBV-monoinfected patients, indicating that low immune pressure is associated with high frequency mutations of the HBV RT region in both serum and CSF. More importantly, most anti-HBV regimens have poor penetration activity through the BBB, which relates to insufficient drug concentrations for suppressing HBV replication. Consequently, subordinate HBV strains harboring resistance mutations in CSF might translocate into peripheral blood and eventually become the dominant HBV strain. For instance, although HBV-DNA was undetectable by qPCR in peripheral blood, the full-length HBV genome and HBV quasispecies carrying LAM resistance mutations were found in CSF samples of HBV-monoinfected patient 3. Our data support the recommendation that a single anti-HBV medicine containing antiretroviral treatment (ART) is not suitable for HIV/HBV-coinfected patients^[28]^.

Our study has some limitations. First, it has been reported that HBV particles can be found in CSF by electron microscopy^[6]^. However, this study did not confirm the existence of HBV viral particles in CSF. Second, the dynamic changes in HBV-DNA levels and quasispecies after antiviral treatment were not observed longitudinally. Third, our study is based on a small-scale population, and some statistics and analysis, such as differences of quasispecies between HIV/HBV coinfected and HBV monoinfected patients, were not performed.

In summary, our study indicates that the CNS is a potential extrahepatic HBV reservoir, which may contribute to the occurrence of HBV drug resistance due to low concentrations of nucleos(t)ide analogs in CSF. Selecting a drug or drug combination with high penetration through the BBB should be considered a strategy for more effective HBV treatment.

## ACKNOWLEDGEMENTS

1. This research was funded by the National Key R&D Program of China (grant numbers 2021YFC2301900-2021YFC2301901).
2. The authors acknowledge the support by Hangzhou Kaitai Biotechnology Co., Ltd (Hangzhou,China).

## REFERENCES

1. Shang Q, Zhang G, Xu C, Chen CX, Yu JG, Yan CY. [Expression of hepatitis B virus antigen in brain tissue from liver cirrhosis patients with hepatitis B and its significance]. Zhonghua Shi Yan He Lin Chuang Bing Du Xue Za Zhi 2001; 15(3):277–279 (in Chinese).

2. Pontisso P, Schiavon E, Alberti A, Realdi G, Locasciulli A, Masera G. Does hepatitis B virus cross the blood‒brain barrier? N Engl J Med 1986; 314(8):518–519.

3. Ene L, Duiculescu D, Tardei G, Ruta S, Smith DM, Mehta S, et al. Hepatitis B virus compartmentalization in the cerebrospinal fluid of HIV-infected patients. Clin Microbiol Infect 2015; 21(4):387 e385–388.

4. Pronier C, Guyader D, Jezequel C, Tattevin P, Thibault V. Contribution of quantitative viral markers to document hepatitis B virus compartmentalization in cerebrospinal fluid during hepatitis B with neuropathies. J Neurovirol 2018; 24(6):769–772.

5. Pettersson JH, Piorkowski G, Mayxay M, Rattanavong S, Vongsouvath M, Davong V, et al. Meta-transcriptomic identification of hepatitis B virus in cerebrospinal fluid in patients with central nervous system disease. Diagn Microbiol Infect Dis 2019; 95(4):114878.

6. Perlejewski K, Bukowska-Ośko I, Rydzanicz M, Pawełczyk A, Caraballo Cortѐs K, Osuch S, et al. Next-generation sequencing in the diagnosis of viral encephalitis: sensitivity and clinical limitations. Sci Rep 2020; 10(1):16173.

7. Nassal M. HBV cccDNA: viral persistence reservoir and key obstacle for a cure of chronic hepatitis B. Gut 2015; 64(12):1972–1984.

8. Chen L, Gan QR, Zhang DQ, Yao LF, Lin RS, Li Q, et al. Increased intrahepatic quasispecies heterogeneity correlates with off-treatment sustained response to nucleos(t)ide analogues in e antigen-positive chronic hepatitis B patients. Clin Microbiol Infect 2016; 22(2):201–207.

9. Coffin CS, Mulrooney-Cousins PM, Peters MG, van Marle G, Roberts JP, Michalak TI, et al. Molecular characterization of intrahepatic and extrahepatic hepatitis B virus (HBV) reservoirs in patients on suppressive antiviral therapy. J Viral Hepat 2011; 18(6):415–423.

10. Best BM, Letendre SL, Koopmans P, Rossi SS, Clifford DB, Collier AC, et al. Low cerebrospinal fluid concentrations of the nucleotide HIV reverse transcriptase inhibitor, tenofovir. J Acquir Immune Defic Syndr 2012; 59(4):376–381.

11. Calcagno A, Di Perri G, Bonora S. Pharmacokinetics and pharmacodynamics of antiretrovirals in the central nervous system. Clin Pharmacokinet 2014; 53(10):891–906.

12. Gunther S, Li BC, Miska S, Kruger DH, Meisel H, Will H. A novel method for efficient amplification of whole hepatitis B virus genomes permits rapid functional analysis and reveals deletion mutants in immunosuppressed patients. J Virol 1995; 69(9):5437–5444.

13. Chen L, Zhang Q, Yu DM, Wan MB, Zhang XX. Early changes of hepatitis B virus quasispecies during lamivudine treatment and the correlation with antiviral efficacy. J Hepatol 2009; 50(5):895–905.

14. Domingo E, Martin V, Perales C, Grande-Pérez A, García-Arriaza J, Arias A. Viruses as quasispecies: biological implications. Curr Top Microbiol Immunol 2006; 299:51–82.

15. Kosakovsky Pond SL, Poon AFY, Velazquez R, Weaver S, Hepler NL, Murrell B, et al. HyPhy 2.5-A Customizable Platform for Evolutionary Hypothesis Testing Using Phylogenies. Mol Biol Evol 2020; 37(1):295–299.

16. Bavaro DF, Calamo A, Lepore L, Fabrizio C, Saracino A, Angarano G, et al. Cerebrospinal fluid compartmentalization of HIV-1 and correlation with plasma viral load and blood-brain barrier damage. Infection 2019; 47(3):441–446.

17. Han Y, Huang LH, Liu CM, Yang S, Li J, Lin ZM, et al. Characterization of hepatitis B virus reverse transcriptase sequences in Chinese treatment naive patients. J Gastroenterol Hepatol 2009; 24(8):1417–1423.

18. Fu Y, Zeng Y, Chen T, Chen H, Lin N, Lin J, et al. Characterization and Clinical Significance of Natural Variability in Hepatitis B Virus Reverse Transcriptase in Treatment-Naive Chinese Patients by Sanger Sequencing and Next-Generation Sequencing. J Clin Microbiol 2019; 57(8).

19. Park ES, Lee AR, Kim DH, Lee JH, Yoo JJ, Ahn SH, et al. Identification of a quadruple mutation that confers tenofovir resistance in chronic hepatitis B patients. J Hepatol 2019; 70(6):1093–1102.

20. Ene L, Duiculescu D, Tardei G, Ruta S, Smith DM, Mehta S, et al. Hepatitis B virus compartmentalization in the cerebrospinal fluid of HIV-infected patients. Clin Microbiol Infect 2015; 21(4):387.e385–388.

21. Inoue J, Ueno Y, Kogure T, Nagasaki F, Kimura O, Obara N, et al. Analysis of the full-length genome of hepatitis B virus in the serum and cerebrospinal fluid of a patient with acute hepatitis B and transverse myelitis. J Clin Virol 2008; 41(4):301–304.

22. Letendre S, Marquie-Beck J, Capparelli E, Best B, Clifford D, Collier AC, et al. Validation of the CNS Penetration-Effectiveness rank for quantifying antiretroviral penetration into the central nervous system. Arch Neurol 2008; 65(1):65–70.

23. Li KW, Kramvis A, Liang S, He X, Chen QY, Wang C, et al. Higher prevalence of cancer related mutations 1762T/1764A and PreS deletions in hepatitis B virus (HBV) isolated from HBV/HIV co-infected compared to HBV-mono-infected Chinese adults. Virus Res 2017; 227:88–95.

24. Chen CH, Hung CH, Lee CM, Hu TH, Wang JH, Wang JC, et al. Pre-S deletion and complex mutations of hepatitis B virus related to advanced liver disease in HBeAg-negative patients. Gastroenterology 2007; 133(5):1466–1474.

25. Wu Y, Zhu Z, Wu J, Bi W, Xu W, Xia X, et al. Evolutionary Analysis of Pre-S/S Mutations in HBeAg-Negative Chronic Hepatitis B With HBsAg < 100 IU/ml. Front Public Health 2021; 9:633792.

26. Schnell G, Price RW, Swanstrom R, Spudich S. Compartmentalization and clonal amplification of HIV-1 variants in the cerebrospinal fluid during primary infection. J Virol 2010; 84(5):2395–2407.

27. Bednar MM, Sturdevant CB, Tompkins LA, Arrildt KT, Dukhovlinova E, Kincer LP, et al. Compartmentalization, Viral Evolution, and Viral Latency of HIV in the CNS. Curr HIV/AIDS Rep 2015; 12(2):262–271.

28. Iser DM, Sasadeusz JJ. Current treatment of HIV/hepatitis B virus coinfection. J Gastroenterol Hepatol 2008; 23(5):699–706.

